# Evolution of personality and locomotory performance traits during a late Pleistocene island colonization in a treefrog

**DOI:** 10.1101/2022.05.15.491936

**Authors:** Roberta Bisconti, Claudio Carere, David Costantini, Anita Liparoto, Andrea Chiocchio, Daniele Canestrelli

## Abstract

Recent empirical and theoretical studies suggest that personality and locomotory performance traits are crucial components of the dispersal syndromes, and that they can evolve during a dispersal process. Island colonisation is one the best characterized processes in dispersal biogeography, and its implication in the evolution of phenotypic traits has been investigated over a wide range of temporal scales. However, the effect of island colonization on personality and performance traits of natural populations has been little explored, and no studies have addressed these processes in the context of late-Pleistocene range expansions. Here, we investigated the contribution of island colonisations triggered by post-glacial range expansions to intraspecific variation in personality and locomotory performance traits. We compared boldness, exploration, jumping performance, and stickiness abilities, in three equidistant populations of the Tyrrhenian tree frog *Hyla sarda*, two from the source area and one from the colonised island. Individuals from the colonised population were significantly bolder than individuals from the source area, as they emerged sooner from a shelter (p=0.028), while individuals from the source area showed markedly higher jumping and stickiness performance (both p<0.001). We discuss these results in the context of the major microevolutionary processes at play during range expansion, including selection, spatial sorting, and founder effects. However, irrespective of the processes contributing the most, our results clearly indicate that late Pleistocene climatic changes have had major consequences not just on species’ range dynamics, but also on the spatial patterns of phenotypic variation within species, including personality and locomotory traits variation.

---

Research on island populations has provided fundamental information on evolutionary diversification and adaptation (Losos and Ricklefs 2009). While it is well established that island populations differ predictably in morphology, size and life history from their mainland counterparts, relatively few studies have documented insularity-driven changes in the behavioural phenotype. A known behavioural feature of island populations of reptiles, birds and mammals is their “tameness”, i.e. the loss of the ability to recognize and respond to predators with competence, a phenomenon that has been associated to relaxed predation pressure and the costs of sustaining the machinery required to detect and escape from predators (Blumstein 2002; Blumstein and Daniel 2005; Cooper et al. 2014; Rödl et al. 2007; Brock et al. 2015; Baeckens and Van Damme 2020). Moreover, in some islands, territoriality is reduced compared to mainland and individuals appear more tolerant towards intruders (Stamps and Buechner 1985; Gray and Hurst 1998). However, in other populations, intraspecific aggression on islands is more intense than on mainland (Vervust et al. 2007; Raia et al. 2010). These patterns are limited to reptiles, birds and mammals and it is unclear why some species/populations “follow the rule” and others less or not, while the mechanisms producing the phenotypic divergence – founder effect, genetic drift, selection, plasticity - are also poorly known (Baeckens and Van Damme 2020). Overall, the picture of the behavioural profiles and responsiveness of island populations is still incomplete.

The study of animal personality, defined as consistent inter-individual variation in sets of behavioural traits with both genetic and epigenetic basis, provides an integrative approach to behaviour (Carere and Maestripieri 2013). Animal personality can influence a wide range of population-level processes with significant ecological and evolutionary implications (Réale et al. 2007; Cote et al. 2010; Wolf and Weissing 2012; Sih et al. 2012; Canestrelli et al. 2016a). In particular, personality traits, such as boldness and exploration have been linked to dispersal, range expansion, and colonization processes in several theoretical and empirical studies (Fraser et al. 2001; Dingemanse et al. 2003; Cote et al. 2010; Chapple et al. 2012; Sih et al. 2012; Canestrelli et al. 2016a; Gruber et al. 2017). Notably, dispersal is a highly energy and resource demanding process that imposes a number of physiological challenges and requires high performance abilities (Phillips et al. 2006; Bonte et al. 2012; Jessop et al. 2018; Kosmala et al. 2017). However, despite the increasing number of studies on personality and dispersal-enhancing traits of animals in many contexts, particularly in invasive species, the role of biogeographic processes in shaping personality traits has not been explored, letting us blind over the temporal dimension of personality traits evolution. It is therefore paramount to delve into the expression of personality and other dispersal-enhancing traits in populations that have colonized islands, taking into account that any suite of traits that enhances departure, arrival and settlement on a given island might, for several reasons (e.g. evolutionary inertia, novel predators and/or competitors), be different from those that allow later generations to successfully survive on that island. For example, information on naturally colonized island populations is virtually absent. A recent study on *Rana temporaria* has actually suggested a differentiation of personality traits in relation to a recent colonization, but the ongoing flow of individuals between mainland and islands could not allow inferences about the causes and the temporal persistence of the observed divergence (Brodin et al. 2013).

Our main goal was to test whether island populations differ from their mainland counterparts in personality and locomotory performance traits, by comparing populations with a known evolutionary history (Foster 1999). We took advantage from the unique opportunity offered by the known colonization history of the Tyrrhenian tree frog (*Hyla sarda)*, a species with an insular distribution in the Western Mediterranean Sea (Bisconti et al. 2011a; Bisconti et al. 2011b). We focused on three equidistant populations, two from Corsica (source) and one from the Elba island (sink, Figure 1), distributed along the route of post-glacial range expansion (as inferred in Bisconti et al. 2011a; Bisconti et al. 2011b; Spadavecchia et al. 2021). The two Corsica populations were sequentially founded via diffusion dispersal, while the Elba population originated via a jump dispersal event (Vences et al. 2003, 2004). Importantly, such expansion processes were recent (post-glacial) and synchronous (Spadavecchia et al. 2021), allowing us to exclude effects due to long-term allopatric divergence. In fact, historical demographic reconstructions showed that this species underwent two sequential expansion steps: the first from Sardinia to southern Corsica, at the end of the Late Plestocene, the second to the remaining part of the current range including northern Corsica and the Tuscan archipelago (Spadavecchia et al. 2021).

**Figure 1.**
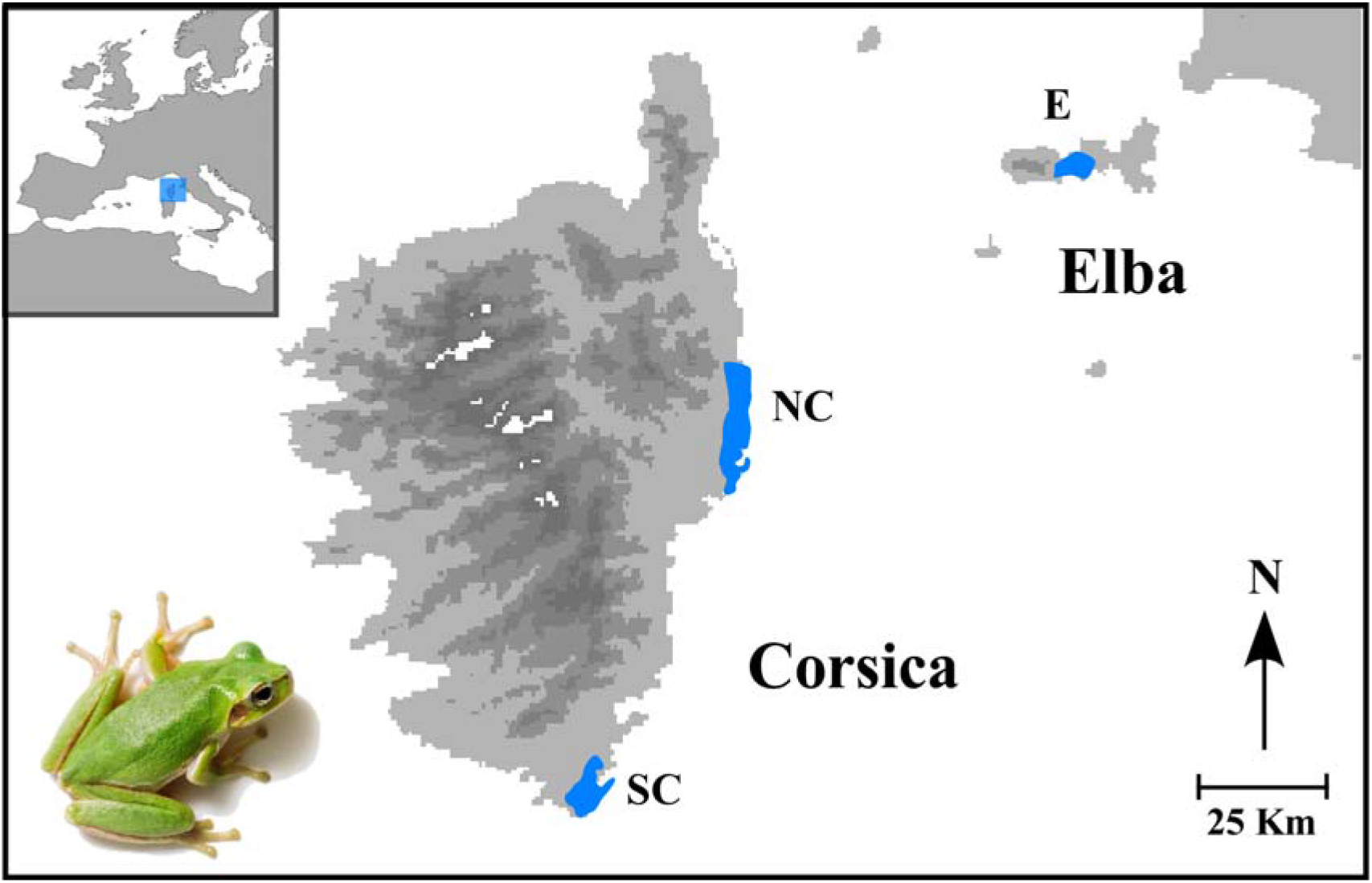
Geographical distribution of the three sampled populations (blue shapes). The study species *Hyla sarda* is shown in the lower left corner.

On the hypothesis that dispersal-related traits have undergone a differentiation in the colonized island, we compared personality (exploration and boldness) and performance (jumping performance at take-off and stickiness) among the three aforementioned populations. Boldness is a personality trait that is associated to both dispersal and risk of predation (e.g., Fraser et al. 2001; Hulthèn et al. 2017). Exploration reflects the individual response to a novelty (*e*.*g*. Verbeek et al. 1994; Cote et al. 2010). Locomotory performance is a key determinant of dispersal ability, and inter-individual variation in jumping performance has been previously analysed in the context of dispersal evolution during (invasive) range expansions (Phillips et al. 2006; Llewelyn et al. 2010; Louppe et al. 2017, Kosmala et al. 2017).

## Material and methods

### Study species

The Tyrrhenian tree frog (*Hyla sarda*) is an amphibian species endemic to the Tyrrhenian islands of Sardinia and Corsica, and the Tuscan archipelago. This arboreal species is rather common at low and intermediate altitudes, with more abundant populations frequently found in coastal areas. (Lanza et al., 2007). Bioclimatic niche models generated for *H. sarda* under both current and periglacial climatic conditions (Bisconti et al. 2011a), showed spatially and temporally homogeneous bioclimatic conditions in most coastal areas, including western Corsica and Elba island. Compared to its closely related species *Hyla arborea*, the Tyrrhenian tree frog is more linked to aquatic habitats, where it breeds in pools and temporary ponds from spring to early summer (Lanza et al. 2007). During Late Pleistocene it colonized the Corsica island from Sardinia taking advantage of a wide and persistent land bridge between the two main islands, and then it reached the Elba island (see Introduction; Bisconti et al. 2011a; Bisconti et al. 2011b; Spadavecchia et al., 2021).

### Sampling and housing

A total of 72 individuals were sampled between April and July 2018 in Corsica (two populations) and Elba island (one population). Following results from previous bioclimatic niche analyses (Bisconti et al., 2011a), all sampling sites (Table 1 and Figure 1) were selected in strictly coastal areas (<10m above sea level, within 1km from the coastline), which allowed us to control for potential effects of bioclimatic differences on the geographic patterns of the investigated traits.

**Table 1.**
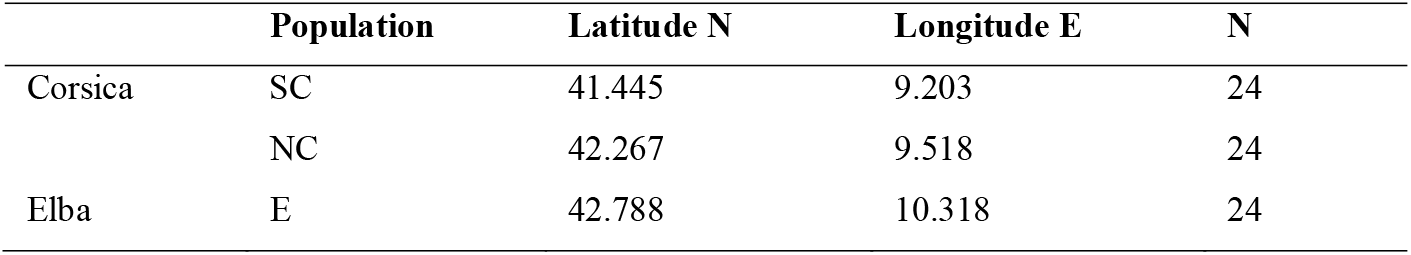
Geographic coordinates and sample size for the three sampled populations of *Hyla sarda*.

|  | Population | Latitude N | Longitude E | N |
| --- | --- | --- | --- | --- |
| Corsica | SC | 41.445 | 9.203 | 24 |
|  | NC | 42.267 | 9.518 | 24 |
| Elba | E | 42.788 | 10.318 | 24 |

Individuals were sampled with hand nets at night after acoustic and visual localization, immediately stored in individual plastic boxes (13,1 cm x 10,2 cm x 4,9 cm) containing humid paper that was renewed daily. Within three days from sampling individuals were transported to our lab facilities, where they were housed under controlled environmental conditions, at a temperature of 24/25 C°, relative humidity of 60-80 % and natural photoperiod. They were kept in individual cages (25 cm x 25 cm x 25 cm) provided with a small dechlorinated water tank (diameter 5 cm), a plant and an oak wood shelter. The cages were not visually or acoustically isolated from each other and were placed randomly with respect to population of origin. Twice a week the cages were cleaned, the water renewed and animals were fed with crickets (*Acheta domestica*). Before starting the tests, the animals were left undisturbed for two weeks except for routine feeding and cleaning duties.

Sampling procedures were performed under the approval of the Institute for Environmental Protection and Research ‘ISPRA’ (protocol # 5944), Ministry of Environment ‘MATTM’ (protocol #8275), Regione Sardegna (#12144) and Corsica (#2A20180206002 and #2B20180206001). Permission to temporarily house amphibians was granted by the Local Health and Veterinary Centre, with license code 050VT427. All handling procedures outlined in the present study were approved by the Ethical Committee of the Tuscia University for the use of live animals. During captivity the animals were monitored daily. No adverse effects on the overall health of tree frogs were observed during the procedures. The animals were released in the original sampling locations at the end of the experimentation.

### Personality traits

For the assessment of personality traits, two tests were performed (Table 2): (i) arena test to investigate the propensity to explore (Figure 2A); (ii) shelter test to examine the boldness-shyness behavioural axis (Figure 2B). Both tests were repeated after 10 days to test for individual consistency over time. All tests were carried out under the same environmental conditions used for housing tree frogs.

**Table 2.**
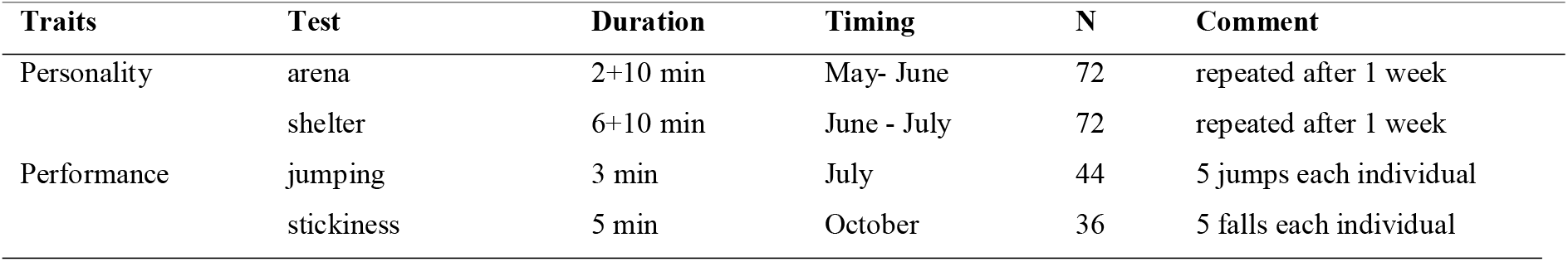
Summary of the tests used to evaluate personality and performance traits. See method section text for details.

| Traits | Test | Duration | Timing | N | Comment |
| --- | --- | --- | --- | --- | --- |
| Personality | arena | 2+10 min | May- June | 72 | repeated after 1 week |
|  | shelter | 6+10 min | June - July | 72 | repeated after 1 week |
| Performance | jumping | 3 min | July | 44 | 5 jumps each individual |
|  | stickiness | 5 min | October | 36 | 5 falls each individual |

**Figure 2.**
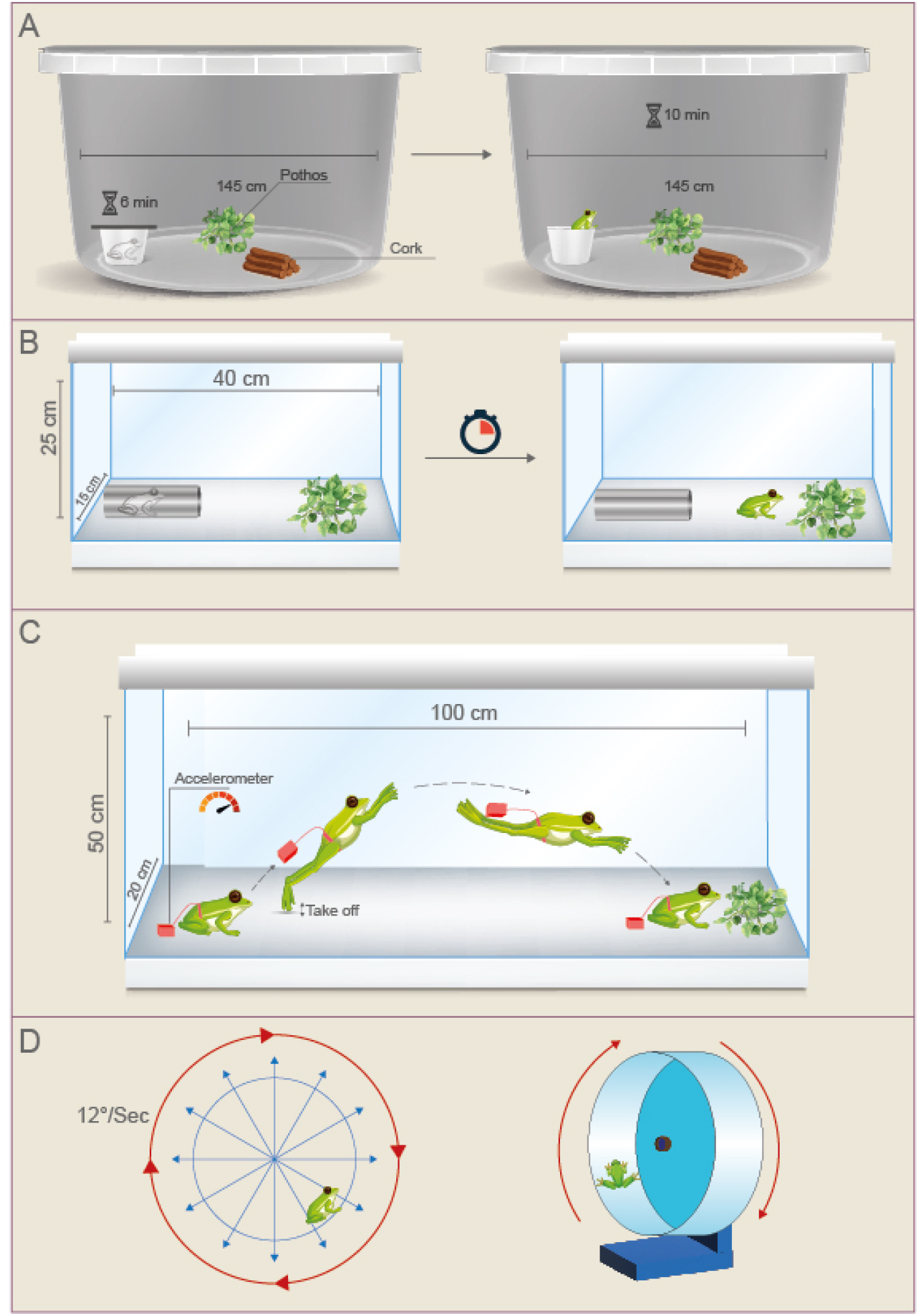
Schematic representation of the tests conducted on *Hyla sarda*. A) Arena test used to investigate the propensity to explore; B) shelter test used to examine the boldness-shyness behavioural axis (as latency to exit from a refugium); C) jumping test to measure the maximum jumping performance at take-off; D) stickiness test used to estimate maximum adhesiveness force.

#### Exploration: arena test

Considering the arboreal life style of tree frogs, we used a cylindric arena with vertical as well as horizontal dimension (diameter 145 cm x height 90 cm) enriched with a plant and a wooden shelter. Individuals were tested randomly respect to population of origin on the same day between 9:00 and 14:00. Each individual was gently introduced in the arena and left undisturbed in a dark jar (5cm x 3 cm) positioned along the border; after two minutes the jar was opened and the animal was allowed to explore the arena for 10 min. After that, it was immediately put back in its home cage. The arena was cleaned after each test in order to remove any chemical cues or secretion left by previous individuals.

The tests were videorecorded (Panasonic DMC-FZ300) and videos were analysed by one operator using Boris 5.1.3 (Friard and Gamba 2016). The following variables were extracted: (i) latency to explore (s); (ii) duration of activity (expressed as percentage on duration of test); (iii) duration of time spent on the arena floor (expressed as percentage on duration of test); (iv) frequency of jumping events.

#### Boldness: shelter test

We used a rectangular arena (length 40 cm x height 25 cm) enriched with a plant. Individuals were tested randomly respect to population of origin on the same day between 9:00 and 14:00. Each individual was gently introduced in a cylindric dark shelter (5cm x 3 cm) positioned along the arena’s border; after 6 minutes the shelter was opened and the animal was given 10 min to exit from the jar. After that, it was immediately put back in its home cage and the arena was cleaned. Tests were video-recorded and videos were visually analysed by one operator to measure the latency to exit from the shelter (s).

### Locomotory performance

For the assessment of performance traits, we carried out two sets of tests (Table 2). Firstly, we measured the maximum jumping performance at take-off (Figure 2C). The jumping force at take-off was measured to characterize the propulsive phase of jump (*e*.*g*. Marsh and John-Alder 1994; Nauwelaerts and Aerts 2006 and reference therein), which is the predominant mode of locomotion in tree frogs. Secondly, we quantified the maximum adhesiveness force using a stickiness test (Figure 2D). Tree frogs have specialised toe pads, which adhere via an area-based wet adhesive mechanism (Emerson and Diehl 1980; Federle et al. 2006; Smith et al. 2006). This anatomical specialization is crucial for a safe landing and is intrinsically linked to jumping performance, as missing the target could have severe consequences in arboreal species, as compared to terrestrial species (Bijma et al. 2016).

All tests were carried out under the same environmental conditions used for housing tree frogs.

#### Jumping test

The jumping tests were conducted in a rectangular arena (length 100 cm x height 50 cm), enriched with a plant placed at the opposite side of the arena, to obtain standardized directionality of tree frog jumps. Individuals were tested randomly respect to the population of origin on the same day between 9:00 and 14:00. The animals were equipped with an accelerometer, deployed by a pelvis-loop harness (Axy-4 units, Technosmart, Rome, 9.15 × 15 × 4 mm, 1 g weight including battery), and induced to jump by a slight stimulation in the pelvic region (Mitchell and Bergmann 2016). The logger was set to record triaxial acceleration (0-4 g) at 100 Hz with 8bit resolution. After that, all individuals were weighted (scale Acculab model ATILON ATL-224-I) and then gently put back in their home cages. The overall vectorial dynamic body acceleration (VeDBA) at take-off was calculated considering a running mean of 30 centi-seconds (Shepard et al. 2008) using Framework 4 software (version 2.5). The highest VeDBA value over five consecutive jumps was used to calculate an individual’s maximum jumping force at take-off (following Shepard et al. 2008).

#### Stickiness test

This test was designed to investigate the degree of adhesiveness, a functional adaptation to the arboreal life style of tree frogs (Duellman and Trueb 1994). We used a transparent rotating wheel (diameter 20 cm x depth 20 cm) with a moderate and constant angular velocity (12°/s). Each individual was gently introduced in a water tank (dechlorinated tap water, at 25°C) for one minute, to standardize hydration level among individuals. Then the individual was transferred to the wheel, and five detachments were collected. We only considered detachments occurring when the head of the animal was oriented in the same direction of the rotation, to differentiate events reflecting maximum stickiness ability of an individual from intentional detachment events. The whole procedure lasted about two minutes per individual. The tests were video recorded (Panasonic DMC-FZ300) and the videos analysed by one operator using the Tracker software (version 4.11.0) to extract the angle of fall (radian). After that, the maximum adhesiveness force was calculated (following Barnes et al. 2006) for all the five detachments performed by each individual, and the best performance was retained for downstream analyses.

### Data analysis

Generalized Linear Mixed effect (GLMM)-based repeatability models (rpt in package ‘rptR’) were used to test the repeatability of each untransformed behavioural traits (Nakagawa and Schielzeth, 2010). In each model, as dependent variable, we entered behavioural traits singly and individuals as random factor. Proportion function (rptProportion in package ‘rptR’) with binomial error distribution and logit link function was used for all the bounded variables (latency, activity, time spent on the floor and latency to exit from shelter), while a Poisson error distribution with log link function (rptPoisson in package ‘rptR’) was applied to count data (occurrence of jumping events; Stoffel et al. 2017). A parametric bootstrapping method (number of iterations=1000) was used to calculate the CI interval and the likelihood ratio test to estimate p-value of the repeatability distribution. Behavioural variables were considered as personality traits and used for the further analysis when their repeatability value was R > 0.2 and its CI excluded zero (Table 3). Preliminary models showed that entering population as fixed factor did not improve the fitting of the model, thus it was no longer considered.

**Table 3.** Summary of GLMM-based repeatability (*R*) estimates from multiplicative model. Parametric bootstrapping (number of iterations=1000) was used to calculate the CI interval and the Likelihood ratio test to estimate p-value of the repeatability distribution. Significant traits are shown in bold.

| Trait | Estimate | 95 % CI | p-value |
| --- | --- | --- | --- |
| Latency to explore | 0.03 | 0-0.074 | 0.149 |
| Activity | 0.03 | 0-0.798 | 0.110 |
| Time on floor | 0.11 | 0-0.254 | 0.107 |
| Jumping events | 0.30 | 0.035-0.522 | **0.013** |
| Latency to exit from shelter | 0.39 | 0.271-0.491 | **<0.0001** |

A second set of analyses was performed to assess the population differences in both personality (boldness and jumping events) and performance traits (jumping and stickiness) by using Linear Models (LM in package ‘lme4’). Dependent variables were log_10_ transformed to meet the assumption of residuals normality. We fit four different models (one for each trait) entering population as fixed factors and checking for all possible contrasts (Table 4). The same approach was used to test whether the same traits differ between the two islands. Preliminary models showed that entering animal body weight as covariate, both with and without interaction with population, did not improve the fitting of the model, thus it was no longer considered.

**Table 4.** Coefficient estimates (±SE) of linear models (LMs) with both personality and performance traits as dependent variable and population as fixed factor. Significant contrasts are shown in bold.

| Dependent variable | Reference Level | Level | Coefficient | SE | t value | Ps(>\|t\|) |
| --- | --- | --- | --- | --- | --- | --- |
| Latency to exit from shelter | NC | SC | -0.071 | 0.287 | -0.246 | 0.807 |
|  | NC | E | -0.576 | 0.287 | -2.003 | **0.049** |
|  | SC | E | -0.505 | 0.273 | -1.852 | 0.069 |
| Jump occurrence | NC | SC | -0.199 | 0.344 | -0.580 | 0.564 |
|  | NC | E | -0.267 | 0.341 | -0.783 | 0.437 |
|  | SC | E | -0.067 | 0.332 | -0.202 | 0.840 |
| Jumping | NC | SC | -1.718 | 1.279 | -1.343 | 0.188 |
|  | NC | E | -8.232 | 1.240 | -6.639 | **0.0001** |
|  | SC | E | 6.513 | 1.184 | 5.503 | **<0.0001** |
| Stickiness | NC | SC | 0.066 | 0.065 | 1.029 | 0.312 |
|  | NC | E | -0.291 | 0.063 | -4.635 | **<0.0001** |
|  | SC | E | 0.357 | 0.055 | 6.518 | **<0.0001** |

Post-hoc comparisons were explored by calculating a standardized effect size Hedges’ (mes in package ‘compute.es’) for the following contrasts: Corsica vs. Elba for both personality and performance traits. The forest-plot (forest in package ‘metafor’) was used to visualize estimates of effect size and their 95% confidence intervals. Effect sizes were considered to be small (Hedges g=0.2, explaining 1% of the variance), intermediate (g=0.5, explaining 9% of the variance), or large (g=0.8, explaining 25% of the variance) according to Cohen (1988).

Finally, we investigated the relationship between personality and performance traits. Generalized Linear Models (GLM in package ‘nlme’) were run entering performance traits (jumping and stickiness) as dependent variable, population as fixed factor and personality traits (shelter and occurrence of jumping events) as covariates. We fitted four different models including all trait combinations. The same approach was used to test the traits relationship between islands.

All statistical analyses were run using R software version 3.5.3 (R Core Team 2019).

## Results

### Repeatability of behavioural traits

GLMMs showed that boldness (r=0.39), measured as the latency to exit from shelter, and the frequency of jumping events during exploration of the arena (r=0.30) were significantly repeatable over time (Figure 3). On the contrary, latency to explore, activity and time spent on the arena floor were not significantly repeatable and therefore they were excluded from the following analyses. The repeatability coefficient with its CI and the p-value of the measured behavioural traits are reported in Table 3.

**Figure 3.**
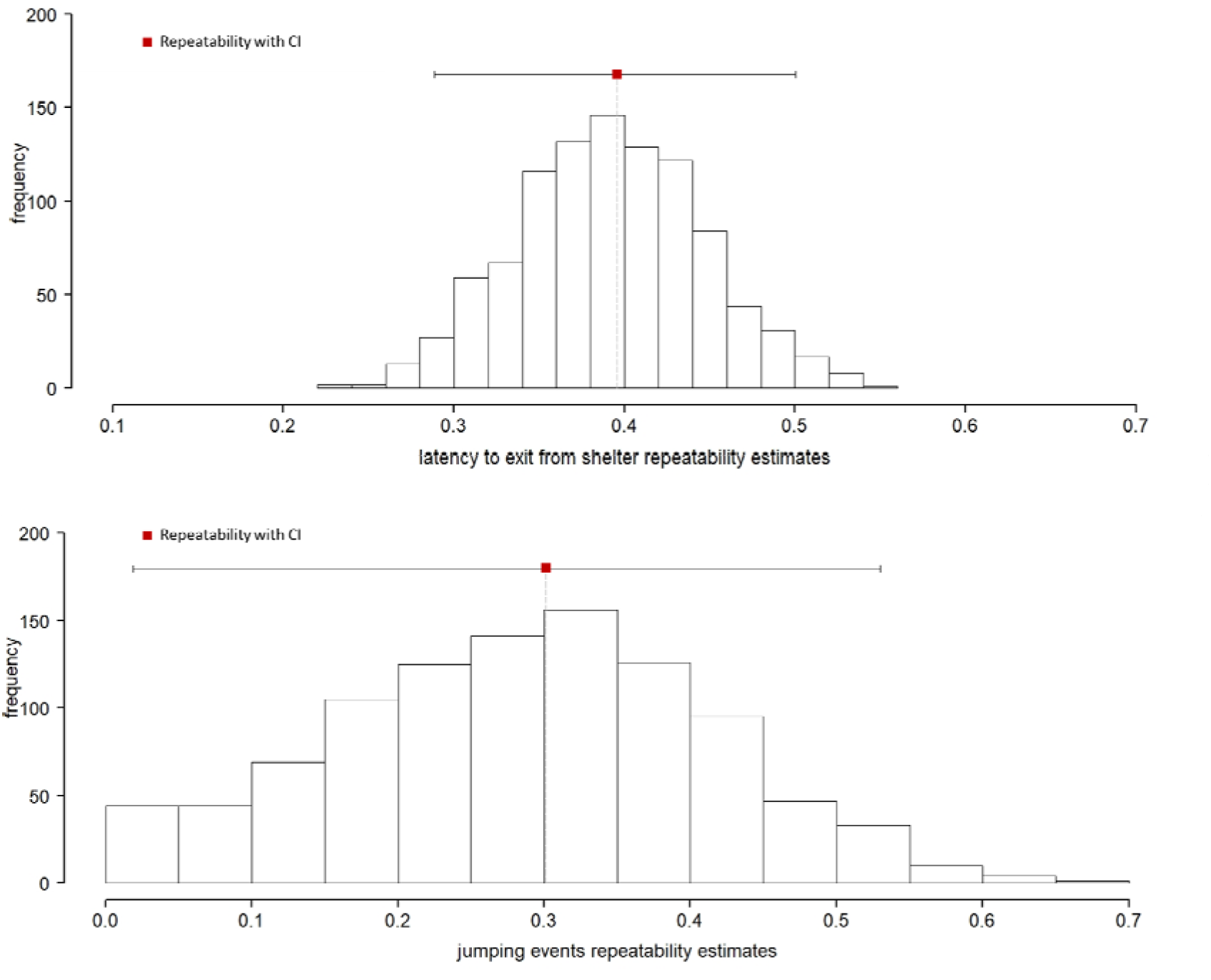
The repeatability estimates: on top latency to exit from shelter of binomial distributed proportion data analysed with logit link; on bottom jumps occurrence of Poisson distributed count data analysed with log link. Link scale repeatability are shown. CI = confidence interval.

### Population differences in personality and performance traits

LMs revealed significant differences in personality traits between north Corsica and Elba populations for boldness (Table 4), which was significantly different also between the two islands pooling together the two Corsican populations (p=0.028). Specifically, tree frogs from Elba island were bolder (emerged sooner from a shelter) than those from Corsica. In fact, the Estimated Marginal Means (EMM) from LMs showed that latency to exit from the shelter was higher in the Corsica populations (SC=4.70 ± 0.193; NC=4.77 ± 0.213) than in Elba (E=4.19 ± 0.193) indicating a more prudent behaviour for Corsica populations (Figure 3). Conversely, jumping frequency during the exploration test did not differ either among populations (Figure 3) as well as between the islands (p=0.575). The parameter estimates of the LM used to test the differences in personality traits among populations are reported in Table 4. Estimate of island’s effect size in boldness, measured as latency to exit from shelter, was large and the 95% CI excludes zero (Figure 4).

**Figure 4.**
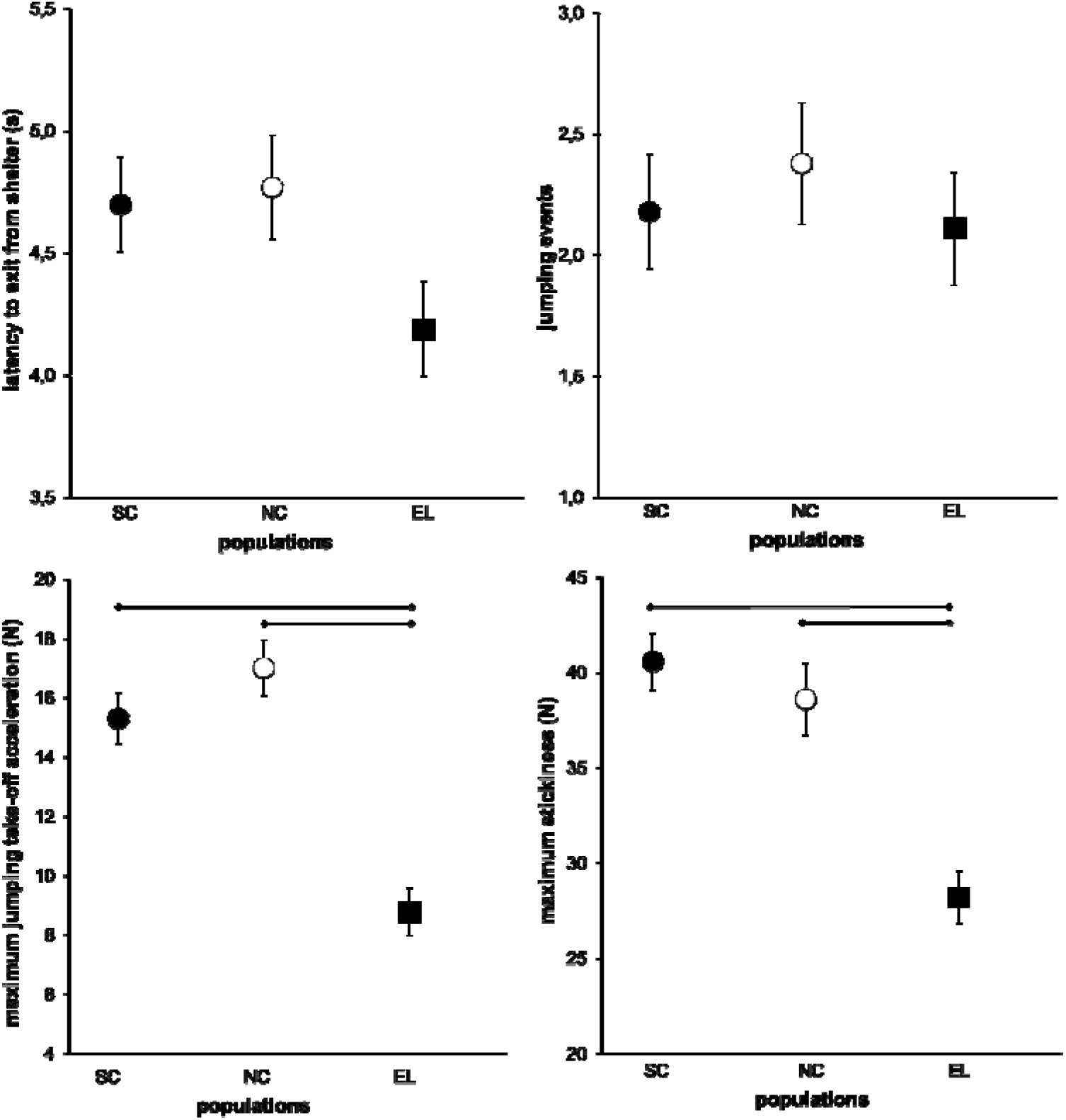
Estimated Marginal Means (EMM) and standard errors from Linear Models (LMs) for personality traits (on top) and performance traits (on bottom). SC= South Corsica (black dot), NC= north Corsica (white dot) and EL=Elba island (black square). Significant contrasts are indicated with bar.

**Figure 5.**
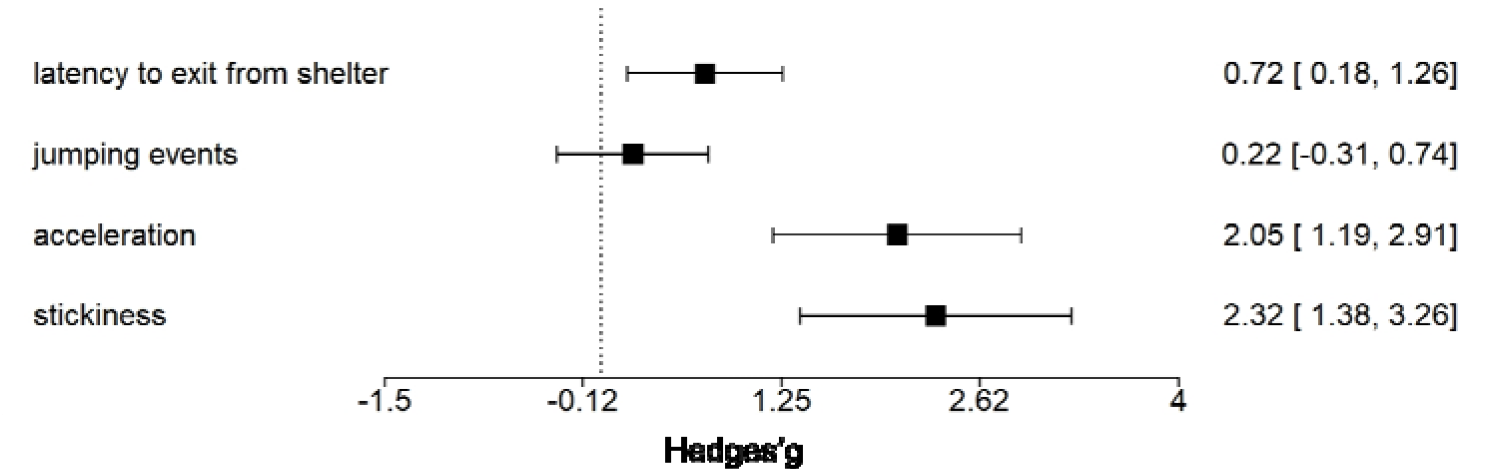
Forest-plot estimating the effect size and 95% confidence interval calculated from all test statistics for both personality and performance traits between islands (Corsica *vs* Elba).

Both performance traits, jumping and stickiness, differed markedly among populations (p<0.0001), except between south Corsica and north Corsica (Table 4). Both Corsica populations showed greater jumping take-off performance than Elba population, as well as higher stickiness. Moreover, both performance traits were significantly different between the two islands (p<0.001). The EMMs values of the maximum jumping take-off performance (N) were twice higher in the north Corsica (NC=17.02 ± 0.942) compared to those of Elba (E=8.79 ± 0.807), while those of the south Corsica lied in the middle (SC=15.31 ± 0.866). Similarly, values of stickiness, measured as the maximum adhesion force (N), were significantly higher in Corsica (SC=40.6 ± 1.48; NC=38.6 ± 1.86) than in Elba (E=28.2±1.36). A summary of the coefficient estimates used to investigate population differences in performance traits is reported in Table 4. Estimate of island’s effect size in performance differences were remarkable and the 95% CI excludes zero for both maximum jumping force at take-off and stickiness (Figure 4).

### Relationship between personality and performance

Significant evidence for association between personality and performance traits, at population level, were found between jumping frequency during the exploration test and the maximum jumping performance. All three populations showed a significant association between these two traits (Table 5).

**Table 5.**
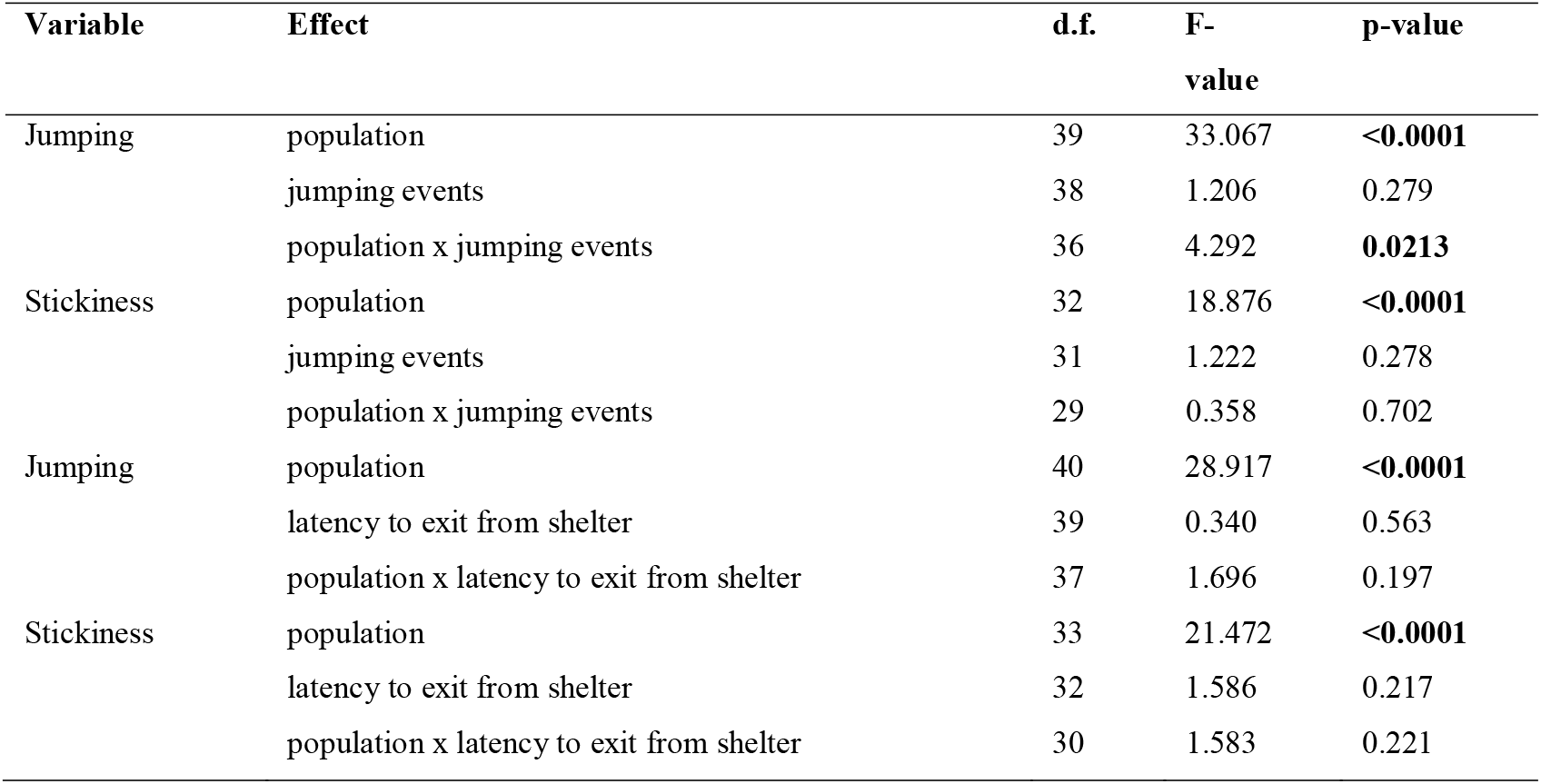
Summary of General Linear Models (GLM) used to validate the association between performance (jumping and stickiness) and personality traits (jumping events and latency to exit from shelter). Significant factors are shown in bold.

| Variable | Effect | d.f. | F-value | p-value |
| --- | --- | --- | --- | --- |
| Jumping | population | 39 | 33.067 | **<0.0001** |
|  | jumping events | 38 | 1.206 | 0.279 |
|  | population x jumping events | 36 | 4.292 | **0.0213** |
| Stickiness | population | 32 | 18.876 | **<0.0001** |
|  | jumping events | 31 | 1.222 | 0.278 |
|  | population x jumping events | 29 | 0.358 | 0.702 |
| Jumping | population | 40 | 28.917 | **<0.0001** |
|  | latency to exit from shelter | 39 | 0.340 | 0.563 |
|  | population x latency to exit from shelter | 37 | 1.696 | 0.197 |
| Stickiness | population | 33 | 21.472 | **<0.0001** |
|  | latency to exit from shelter | 32 | 1.586 | 0.217 |
|  | population x latency to exit from shelter | 30 | 1.583 | 0.221 |

## Discussion

The present study provides evidence of a substantial differentiation in personality and locomotory performance traits of a tree frog population residing on a recently colonised island (Late Pleistocene). We found that tree frogs from Elba island (sink) were significantly bolder than their ancestors (Corsica), which suggests a less prudent behaviour. Moreover, individuals from Elba island showed a markedly lower performance than those from Corsica. To our knowledge, this is the first study to provide empirical data relating the evolution of personality and dispersal-related traits to past biogeographic events, as previously just hypothesised (Canestrelli et al. 2016a). Such significant differentiation could not be explained by long-term allopatric divergence due to the isolation of Elba from Corsica populations, since Elba island and the north of Corsica were colonised as the last step of a single and recent range expansion event (less than 10ka; Spadavecchia et al., 2021). While this island colonisation event likely triggered the observed phenotypic differentiation, it might imply a number of different eco-evolutionary processes at play during and after the colonization, moulding spatial patterns of behavioural and performance traits, as detailed below.

The first one is selection in favour of dispersal enhancing traits by spatial sorting (Shine et al., 2011). This process has been invoked to explain changes in dispersal-related phenotypes along an expansion route, in particular at the range front of an expanding population (Lowe et al. 2015; Canestrelli et al. 2016b; Phillips & Perkins 2019). Mounting evidence on invasive species suggests that spatial sorting processes could promote directional changes in the phenotypic and genotypic make-up of populations during range expansions (e.g. Travis & Dytham 2002; Phillips et al. 2010; Shine et al. 2011; Brown et al. 2014). In our case, with a spatial sorting process at play, phenotypic differences should have been observed between southern and northern Corsica populations as well. Instead, the lack of differentiation between the two Corsica populations (Figure 3 and Figure 4), together with the temporal synchrony of the colonization events of northern Corsica and Elba island, make the role of spatial sorting in explaining the observed pattern very unlikely.

Another hypothesis is the possible role played by ecological release in Elba island. This process is the expansion of range, habitat and/or resource use by an organism after arrival in a new community, it is often invoked for populations upon initial island colonization (Gillespie 2009). Ecological release may have caused a shift in the behavioural repertoire of the founding population, leading to an increase in the expression of boldness (Blondel, 2000; Novosolov et al. 2013; Baeckens and Van Damme 2020). However, Elba is a continental island, with ecological communities that do not substantially differentiate from mainland and, to the best of our knowledge, it did not undergo any massive change in animal and plant communities in the recent past. Despite we can’t fully exclude a partial role for this process, we see no evidence supporting it.

By controlling important microhabitat variables, bioclimatic differences between islands might have promoted the observed differentiation upon colonization of the Elba island (e.g. Martìn et al., 2005). However, results from a bioclimatic niche modelling clearly showed a comparable situation between the two islands, leading us to exclude a role of past/present climatic conditions in explaining such pattern of differentiation (Bisconti et al., 2011b). Likewise, we could also exclude any latitudinal effect that should have caused a differentiation between southern and northern Corsica, and the Elba island.

The most plausible scenario that could explain the origin of the observed differences is a founder event associated to the colonization of the Elba island. The lower genetic diversity of the Elba island population as compared to the rest of the species’ range (Bisconti et al. 2011a), is a typical outcome of genetic drift following island colonization (Wright 1931; Waters et al. 2013), and strongly supports a scenario of phenotypic divergence via founder event upon colonization. Indeed, this study shows a substantial divergence in personality and performance traits in an isolated environment compared to the source population. At the same time, despite this is a parsimonious explanation of the observed pattern, it does not rule out a role also of environmental filtering exerted by island recurring conditions or other factors as well. For example, since dispersal-related of phenotypic traits might be affected by land-cover features (Shine et al., 2021; Brown et al., 2006), and vegetation patterns are plausibly relevant for a partially arboreal species like *H. sarda*, a comparative and retrospective analysis of the past and current vegetation coverage of the two islands Corsica and Elba could contribute to a better explanation the observed differentiation.

Finally, we would like to emphasize that, to the best of our knowledge, our study is one to the few so far showing repeatable inter-individual differences in jumping frequency, a trait plausibly related to locomotory performance in an insular context. Jumping frequency in tree frogs could relate to predator-escape ability, a trait which would be therefore under selection, as hypothesized for sprint speed in lizards (Hoskins et al., 2017).

The study of island populations will provide fundamental insights in the evolution of personality, as it did for the other, more studied aspects of the individual phenotype. Studies on aggressive and acoustic behaviour as well as on flight distance, indicator of risk-taking behaviour (e.g. Møller 2008, Hamao et al. 2021), and other specific anti-predatory responses on this and other island species and populations, together with an analysis of local ecological and demographic factors, especially predation pressure and population density, should be the following steps to achieve a more complete picture of the causes of the peculiar behavioural phenotype of island populations.

## Author Contributions

D.Can., C.C., and R.B. designed research; D.Can., R.B., A.C., and A.L. performed research; A.L., D.Cos. and C.C. analyzed data, R.B. and C.C. wrote the paper with inputs from the other authors.

## Acknowledgments

We warmly thank all the collaborators that assisted in sampling and housing: Armando Macali, Alessandro Carlini, Cinzia Mastrogiovanni, Ambra Pazzani, Francesco Ciabattoni and Lorenzo Latini. We also thank Giacomo Dell’omo and the entire staff of Technosmart for having provided the accelerometer and the kindly advise on its use.

## Funding

This work was supported by a grant from the Italian Ministry of Education, University and Research (PRIN project 2017KLZ3MA).

## Conflict of Interest

No conflict of interests.

## Notes

### Competing Interest Statement

The authors have declared no competing interest.

